# Markov Spatial Flows in Bold Fmri: A Novel Lens on the Bold Signal Reveals Attracting Patterns of Signal Intensity

**DOI:** 10.1101/2023.12.15.571860

**Authors:** Robyn L. Miller, Victor M. Vergara, Erik B. Erhardt, Vince D. Calhoun

## Abstract

While the analysis of temporal signal fluctuations and cofluctuations has long been a fixture of blood oxygenation-level dependent (BOLD) functional magnetic resonance imaging (fMRI) research, the role and implications of spatial propagation within the 4D neurovascular BOLD signal has been almost entirely neglected. As part of a larger research program aimed at capturing and analyzing spatially propagative dynamics in BOLD fMRI, we report here a method that exposes large-scale functional attractors of spatial flows formulated as Markov processes defined at the voxel level. The brainwide stationary distributions of these voxel-level Markov processes represent patterns of signal accumulation toward which we find evidence that the brain exerts a probabilistic propagative undertow. These probabilistic propagative attractors are spatially structured and organized interpretably over functional regions. They also differ significantly between schizophrenia patients and controls.

## 1. INTRODUCTION

The patterns of activated brain space measured by BOLD fMRI are typically investigated via pairwise correlations between timeseries corresponding to a fixed collection of functionally-identifiable brain regions or distributed networks [1-3]. This reduced representation of the BOLD signal is referred to as static functional network connectivity (sFNC) when computed over the full recording, and as dynamic functional network connectivity (dFNC) when correlations are evaluated on successive sliding windows through the scan. Dynamic functional network connectivity can capture large scale patterns of brainwide coactivation that change with time, but this common paradigm yields no information about explicitly spatial signal flows, nor about the role of such flows in supporting or impeding the nodal connectivity patterns commonly associated with healthy brain function. Addressing this persistent blind spot in fMRI analyses opens new avenues for gaining scientific and clinical insight from BOLD fMRI.

## 2. METHODS

### 2.1 Data and Group ICA

We use data from a large, eyes-closed resting-state functional magnetic resonance imaging (fMRI) study with approximately equal numbers of schizophrenia patients (SZs) and healthy controls (HCs) (*n*=314, nSZ=151). All subjects in the study signed informed consent forms. Scans were preprocessed according to a standard, previously published pipeline [4] and decomposed with group independent component analysis (GICA) into a set of 100 group-level spatially independent component spatial maps (SMs) and corresponding subject-specific timecourses. Through a combination of automated and manual pruning, *N*=47 (**Figure 1**) functionally identifiable networks are retained: 5 subcortical (SC) networks; 2 auditory (AUD) networks; 11 visual (VIS) networks; 6 sensorimotor (SM) networks; 13 cognitive control (CC) networks; 8 default mode (DMN) networks; 2 cerebellar (CB) networks.

**Figure 1.**
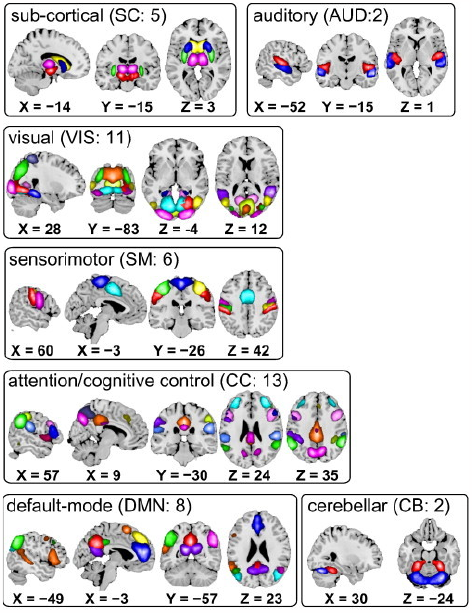
Composite maps of the 47 functional networks, sorted into seven functional domains. Each color in the composite maps corresponds to a different network [4]

**Figure 2.**
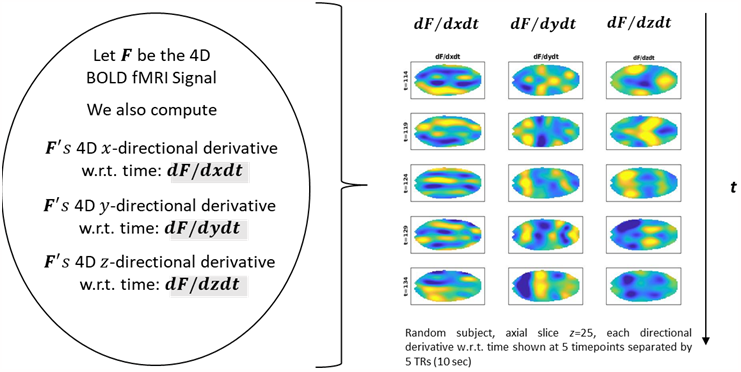
Local directional flows. Space-time directional derivatives computed at voxel and each timepoint using the central difference method. Examples (right) of evolving *x*-direction (column 1), *y*-direction (column 2) and *z*-direction (column 3) space-time derivatives.

**Figure 3.**
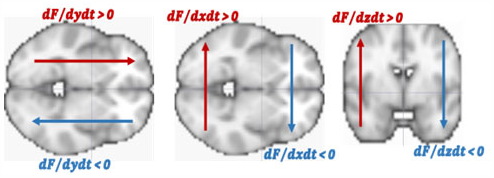
Signed directions along each dimension. displayed on axial and coronal brain slices: dF/dxdt (left), F/dydt (middle), dF/dzdt (right) with positive direction along each dimension in red, negative direction in each direction in blue.

**Figure 4.**
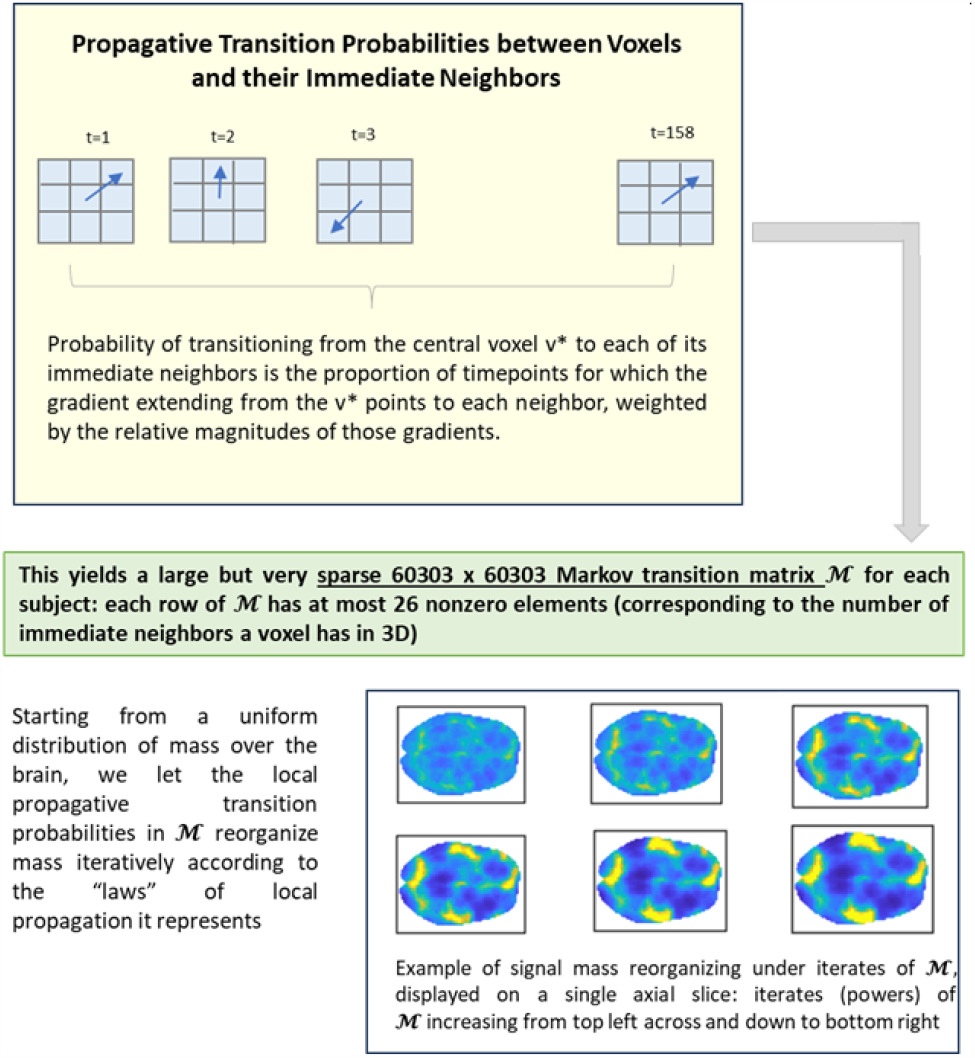
Constructing and running the gradient-driven Markov process: (Top) Planar schematic showing local space-time gradients inducing nearest-neighbor transition probabilities; (Bottom) Example from one subject showing powers of the subject’s Markov matrix successively reorganizing signal mass in a planar axial slice.

### 2.2 Data and Spatiotemporal Gradients (STGs)

Additional processing was performed on scans prior to computing directional spacetime derivatives. This second-stage processing consisted of smoothing preprocessed scans with a 3D Gaussian kernel (*σ* = 3) and 1D temporal moving average with windows of length 3. The gray matter mask for this data contained 60303 voxels: the *x*-dimension (coronal) has length 53; the *y*-dimension (sagittal) has length 63; the *z*-dimension (axial) has length 46. There are 158 sampled timepoints in each scan. To diminish confounding of spatiotemporal gradients by subject differences in global signal amplitude, each fully processed fMRI volume is rescaled by its own mean amplitude. For each voxel *v* = (*x, y, z*) ∈ ℤ^53X63X46^ and each TR *t* ∈ {1,2, …,158} in an fMRI volume, let ***F***(*v, t*) = ***F***(*x, y, z, t*) denote the amplitude-normalized fMRI signal at (*x, y, z, t*). At every voxel and timepoint, we now consider 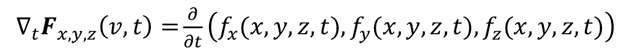 where: *f*_x_ = *d****F***/*dx, f*_y_ = *d****F***/*dy* and *f*_z_ = *d****F***/*dz*. The numerical directional and space-time derivatives are computed via the central difference method, e.g.,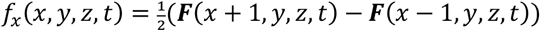 and 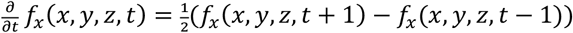.

This yields three new volumes 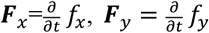 and 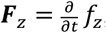, each of the same dimensionality as ***F***.

### 2.3 Gradient-Driven 3D Markov Flow in the Brain *Nearest-Neighbor Markov Process*

From each voxel *v*^*^ = (*x*^*^, *y*^*^, *z*^*^) there is unique vector 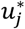 joining the central point in *v*^*^ to the centers of each of its c = 1,2, …, *k*^*^ ≤ 26 immediate neighbors *n*_j_(*v*^*^). In what follows we replace 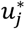 with its unit-normalization 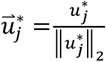. The dot product of the unit normalized space-time gradient 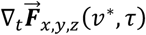 has maximum positive dot product with one of these normalized “spokes, 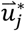, suggesting flow at *v*^*^ is moving more toward the corresponding neighbor, *n*_j_(*v*^*^), than other neighboring voxels at time τ. The evidence for the flow through *v*^*^ at time τ favoring target voxel *n*_i_(*v*^*^) is stronger when the gradient magnitude *w*^*^(τ) = ||∇_t_***F***_x,y,z_(*v*^*^,τ)||_2_ is larger, weaker when the gradient magnitude *m*^*^(τ) is smaller. Let 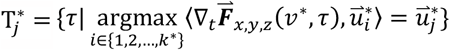 be the set of timepoints at which flow through *v*^*^ is most directionally aligned toward neighbor *n*_j_(*v*^*^). Now sdefine the *gradient-driven transition probability*, 𝒫 (*n*_j_(*v*^*^)| *v*^*^) between a voxel *v*^*^ and its *J*^th^ immediate neighbor *n*_j_(*v*^*^) as

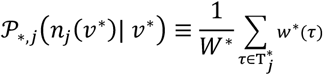

where 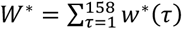. In words, the probability that signal “mass” at voxel *v*^*^ flows into each of its immediate neighbors *n*_i_(*v*^*^), *i* = 1,2, …, *k*^*^ ≤ 26 is simply the summed magnitudes of gradients anchored at *v*^*^ pointed at that specific neighbor, rescaled by the sum of all gradient magnitudes originating at *v*^*^. This construction yields a large, sparse 60303 x 60303 Markov matrix:

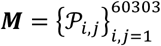

in which every row sums to one and contains at most 26 non-zero elements

#### Stationary Distributions

In practice, all the Markov processes thus constructed from our data have been ergodic (both irreducible and aperiodic), ensuring the existence of a limiting stationary distribution: i.e., a 60303-element distribution π such that π***M*** = π, which was always realizable as *μ****M***^N^ for *N* < 200 in our data, where *μ* is the uniform distribution on {1,2, …,60303} (see **Figure 6**). For brevity, going forward we will refer to these stationary distributions as *Markov propagative attractors* (MPAs).

#### Functionally-Localized Markov Propagative Attractors (fMPAs)

The effect of applying powers of ***M*** to a voxelwise uniform initial condition can also be functionally-localized (see **Figure 5** and **Figure 6**) by averaging over voxels within each functional network mask. The mask for network *i* contains the upper 5% of voxels in its z-scored spatial map. Any voxel in the upper 5^th^ percentile of more than one component map is assigned to the network in which its value is largest. A functionally-localized version of the fMRI signal itself (fBOLD) is computed simularly, averaging the signal within each network mask at each TR.

**Figure 5.**
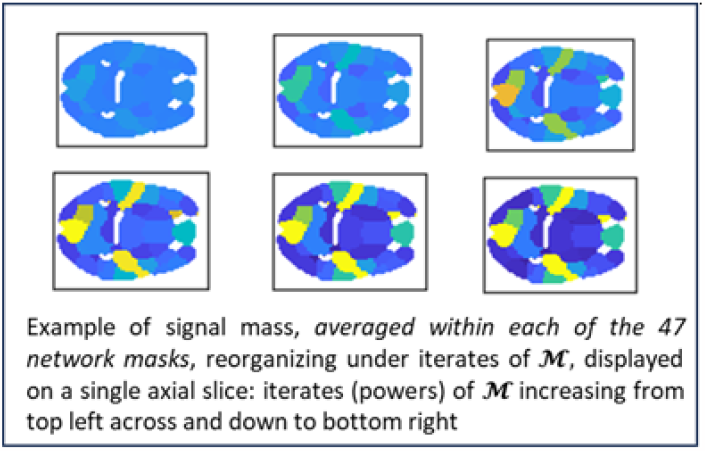
Functionally-localized. signal reorganization under powers of voxel-level nearest-neighbor Markov matrix *M*. Example shows powers of one subject’s Markov matrix successively reorganizing network-averaged signal mass in a planar axial slice.

**Figure 6.**
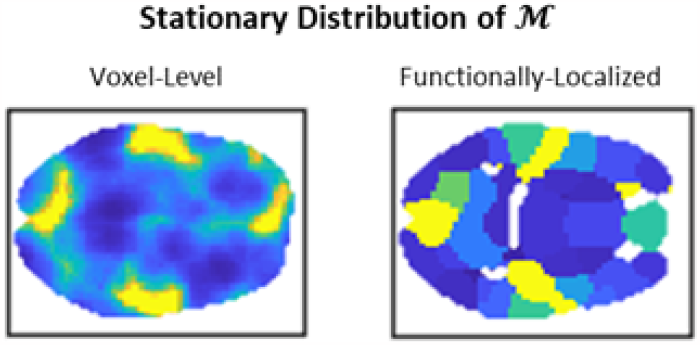
Stationary distribution of the Markov matrix ***M*** displayed on an axial slice; (Left) voxel-level stationary distribution; (Right) Functionally-localized version of the stationary distribution.

#### Imprint of Static fMPAs in the Dynamic fMRI Signal

The whole brain MPA and functionally localized fMPAs and both compact representations of the local flow dynamics captured in the 60303x60303 Markov matrix ***M***, the unique limiting distribution of signal mass induced by the 3D nearest-neghbor Markov process built from local gradients in each subject’s scan. The distribution need not ever be realized during a scanning session and the highly fluid dynamic fMRI signal will at best carry weak, transient evidence of this theoretical object. To capture the strength of these static theoretical distributions in an empircal time varying signals we developed a *dynamic imprint* metric (see **Figure 7**), ***d****𝒥*, defined from the spatial correlation between the fixed fMPA and the dynamic fBOLD signal at each TR as the proportion of TRs for which the Pearson correlation between the fMPA and fBOLD at t is *both positive and significant (p<0*.*05)*:

**Figure 7.**
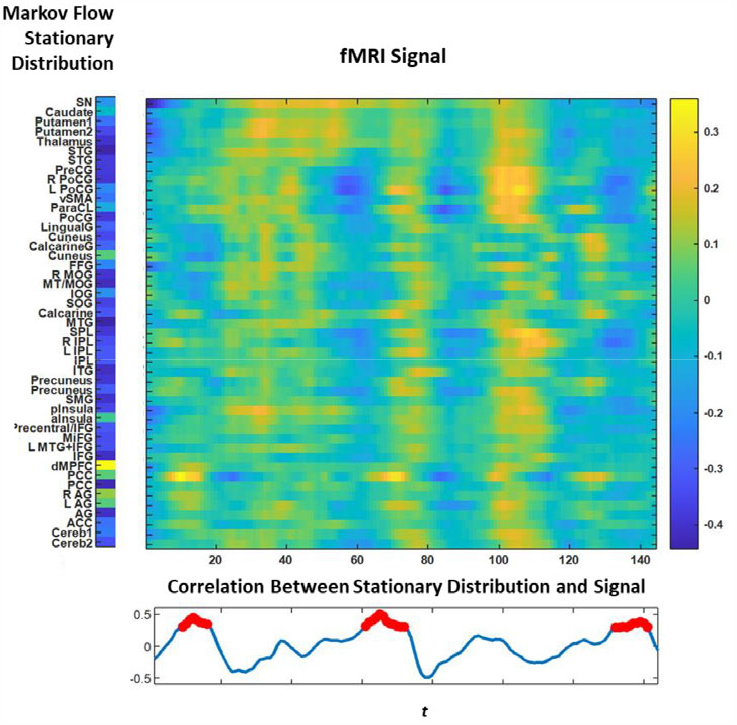
Schematic showing ingredients of the dynamic imprint metric between a fMPA and the functionally localized fBOLD signal: (left) static fMPA; (middle top) dynamic fBOLD; (middle bottom) time varying spatial correlation between fMPA and fBOLD with timepoints at which the correlation is positive with p-value less than 0.05 marked in red; ***d*** 𝒥 is the proportion of such points, in this case ***d*** 𝒥 = 0.2.

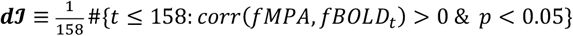

### 2.4 Statistical Analysis

The SZ effects reported here are obtained through a multiple regression model on SZ diagnosis, gender, age and motion. Displayed effects are significant (*p* < 0.05) after correction for multiple comparisons, unless otherwise indicated. Colormaps are symmetric about zero, with cooler (resp. warmer) colors indicating negative (resp. positive) SZ effects.

## 3. RESULTS

The functionally-localized stationary distributions that arise from the gradient-driven nearest-neighbor Markov process detailed above offer a different lens on the neurovascular BOLD signal, capturing global functional implications of directional flow probabilities specified at the voxel level. We find, broadly, that these flows move signal mass away from higher cortical regions such as the default mode and cognitive control networks toward sensory and subcortical areas. The cingulate cortices (posterior (PCC) and anterior (ACC) of the default mode network are exceptions to this general trend (). There are notable differences between the patient and control groups in which areas are significantly gaining or losing signal mass (and how much) under the Markov flow (see **Figure 8** and **Figure 9**). In particular, we see that for SZ patients in contrast with controls, certain default mode regions are lower order “donors” of signal mass to the rest of the brain (e.g. the angular gyri (LAG, AG, RAG)) (see **Figure 8** and **Figure 9**), while others become net beneficiaries of flow from other regions of the brain (e.g. dorsolateral prefrontal cortex (dMPFC)) (again, **Figure 8** and **Figure 9**).

**Figure 8.**
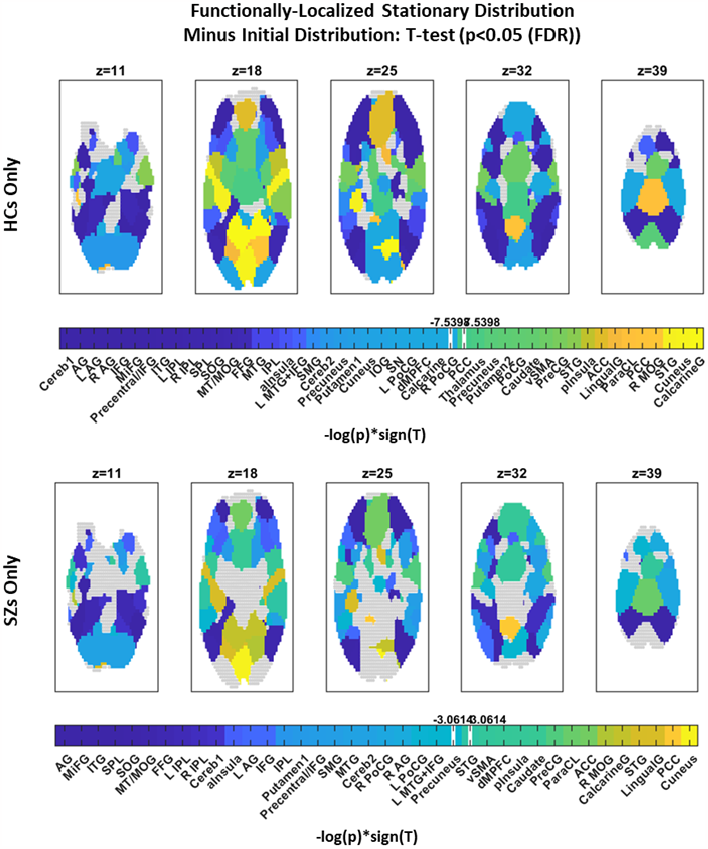
Significant T-tests for difference between functionally-localized stationary distribution and initial uniform condition displayed in select axial slices: (Top) Healthy controls only; (Bottom) SZ patients only; cold colors mark regions that have lost signal mass under the Markov process; warmer colors are regions that have gained signal mass. Colorbars indicate FDR bounds with numerically labeled white stripes. The Markov matrix is volume-preserving, so starting from a uniform initial distribution means that unit volume is being redistributed under powers of ***M*** but the total mass is always one, i.e. anything gained by one region, had to be relinquished by another. Note that more regions exhibit significant T-stats for controls than for patients. Also note that for SZs, dMPFC gains signal mass under powers of ***M***.

**Figure 9.**
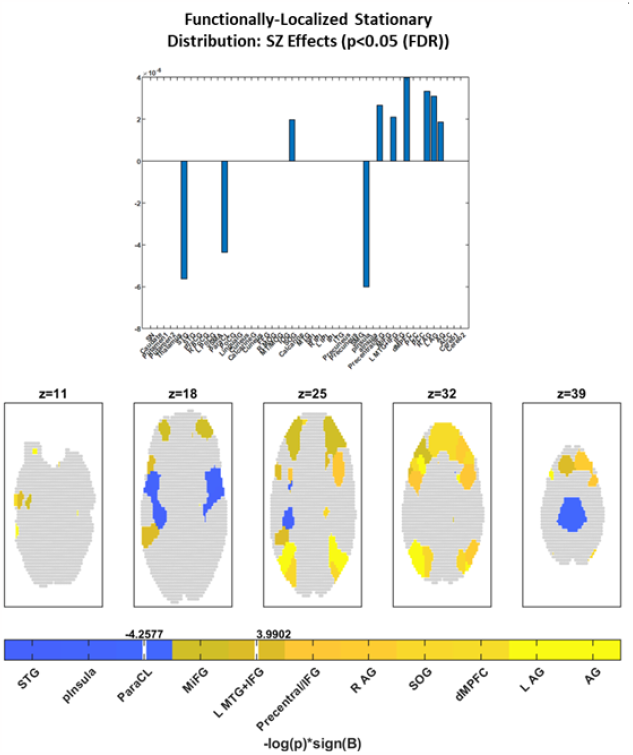
(Top) Significant (FDR) SZ regression effects on network-level stationary distributions and the uniform initial in select axial slices; (Bottom) Significant SZ regression effects on functionally-localized stationary distribution displayed in select axial slices: cold colors indicate regions where the stationary distribution is lower in SZs; warmer colors are regions where the stationary distribution is higher in SZs. Colorbar indicates FDR bounds with numerically labeled white stripes (all networks corresponding to colors warmer (resp. cooler) than the positive (resp. negative) bound remain significant after FDR correction). SZ exhibits positive effects on several default mode networks: e.g. angular gyri and dorsomedial prefrontal cortex.

Moreover, we find evidence tha 1.t subject-specific fMPAs present significantly greater dynamic imprint, ***d*** 𝒥 in the subject’s fBOLD signal than does the population mean fMPA (T-test for ***d***𝒥 (subject fMPA, subject fBOLD) vs. ***d***𝒥 (pop avg. fMPA, subject fBOLD) yields *T* = 3.02, *p* = 0.002); 2. the dynamic imprint in a subject’s fBOLD signal of the mean fMPA for healthy controls only is significantly smaller in patients than in controls: (regression of ***d***𝒥 (healthy mean fMPA, subject fBOLD) on age, gender, motion and SZ diagnosis yields a negative SZ effect with *p* = 1.12*e* - 06), suggesting that the functionally-localized Markov stationary distributions – a theoretical corollary of their voxel-level nearest neighbor Markov flows – derived from healthy individuals are much more weakly present in patients with schizophrenia.

## 4. DISCUSSION

We present here a novel way of approaching the BOLD fMRI signal, focusing on functional reorganization of signal intensity induced by Markov processes built from highly local probabilistic trends in directional flow specified at the voxel level. The preliminary findings suggest that a spatial undertow in the BOLD signal is weakly pulling energy *toward* cingulate cortices (PCC and ACC), *away from* other regions in the default mode and executive/cognitive control networks and broadly *toward* subcortical and sensory areas. SZ significantly elevates mass in several default mode areas, including angular gyri and dorsomedial prefrontal cortex (dMPFC); posterior insula loses more mass under the nearest-neighbor Markov flow in SZ patients than controls. Schizophrenia is a complex mental illness with multiple causal pathways that resting state BOLD fMRI has only begun to elucidate. In this highly preliminary work, we demonstrate one potential role for local spatial flows in characterizing how the brains of schizophrenia patients, as reflected in the BOLD signal, differ from those of healthy controls.

These stationary distributions also present evidence of being more present in subject scans than a generic, biologically plausible distribution (in this case, simply the population average fMPA), suggesting some relevance of a theoretical, not necessarily realized, byproduct of instantaneous voxel-level directional propagation to the observed BOLD signal.

This work is still in preliminary stages, with many weaknesses remaining to be addressed. There are questions of robustness to smoothing parameters, the optimal time window on which to compute transition probabilities (with 26 nearest neighbors, the interval should be long enough to capture more than one transition to each, but we also may want to factor in temporal non-stationarities in the Markov matrix). In this initial foray into a Markovian formulation of local BOLD spatial flows, we have weighted the transition probabilities with gradient magnitudes but have not, e.g. considered scenarios in which small enough magnitudes provoke no transition or large enough magnitudes project transitions beyond the radius of immediate neighbors. All of this to say that the work reported here is preliminary and exploratory, but also highly novel and innovative with respect to currently dominant paradigms of fMRI analysis. We hope that this work motivates further development of approaches to BOLD fMRI that focus explicitly on spatially propagative factors in the signal.

## 5. ACKNOWLEDGEMENTS

This work was supported by NIH R01MH123610 and NSF 2112455.

